# Cytologic, Genetic, and Proteomic Analysis of a Yellow Leaf Mutant of Sesame (*Sesamum indicum* L.), *Siyl-1*

**DOI:** 10.1101/449967

**Authors:** Tongmei Gao, Shuangling Wei, Jing Chen, Yin Wu, Feng Li, Libin Wei, Chun Li, Yanjuan Zeng, Yuan Tian, Dongyong Wang, Haiyang Zhang

## Abstract

Leaf color mutation in sesame always affects the growth and development of plantlets, and their yield. To clarify the mechanisms underlying leaf color regulation in sesame, we analyzed a yellow-green leaf mutant. Genetic analysis of the mutant selfing revealed 3 phenotypes—*YY*, light-yellow (lethal); *Yy*, yellow-green; and *yy*, normal green—controlled by an incompletely dominant nuclear gene, *Siyl-1*. In *YY* and *Yy*, the number and morphological structure of the chloroplast changed evidently, with disordered inner matter, and significantly decreased chlorophyll content. To explore the regulation mechanism of leaf color mutation, the proteins expressed among *YY*, *Yy*, and *yy* were analyzed. All 98 differentially expressed proteins (DEPs) were classified into 5 functional groups, in which photosynthesis and energy metabolism (82.7%) occupied a dominant position. Our findings provide the basis for further molecular mechanism and biochemical effect analysis of yellow leaf mutants in plants.

## Introduction

Sesame (*Sesamum indicum* L.) is a valuable crop with 45–63% oil in its decorticated seeds, which also contain numerous beneficial minerals, antioxidants, and multi-vitamins [1]. Compared with other oilseed crops, sesame is still a low yield crop with low harvest index. The key research objectives in sesame are to increase the photosynthesis efficiency and the yield per square area.

Leaf color is an important trait related with the chlorophyll content, and always affects the photosynthesis efficiency and the final productivity [2]. In plants, the chloroplast structure always alters the composition and content of photosynthetic pigments. Several studies on the leaf color mutants have uncovered a wide range of leaf color mutation types [3]. Most mutagenesis of leaf color were related to the structure and function of the chloroplast [4], chlorophyll biosynthesis and degradation mechanisms [5], photosynthesis [6], chloroplast developmental characteristics [7], genetics [8], and molecular mechanisms [9]. Furthermore, some mutants were used for crop breeding to higher biomass, quality, and/or other specific characteristics [10]. There is a rich diversity of leaf color mutation types in plants [11–14]. Leaf color mutants were divided into five types according to the color classification albino, yellowing, light-green, stripe, spot, etc [15, 16]. Falbel [12] divided the chlorophyll mutant into two categories: (1) mutants that lack chlorophyll b, such as *arabidopsis* mutants [17], and (2) mutants with reduced synthesis of total chlorophyll and chlorophyll b; currently most mutants belong to the latter category. Leaf color characteristics controlled by both nuclear inheritance and cytoplasmic inheritance, may be a quantitative or a quality trait [18]. Most of the leaf color variations are caused by nuclear gene mutation inducing a type of chlorophyll deficiency in higher plants. The leaf color variations are mostly monogenic mutation, while a handful of them are polygenic mutation. Among these mutations, the recessive mutations are predominant and the number of dominant mutations are few [2, 19]. Cytoplasmic inheritance accounts for a small proportion of the leaf color mutants [20]. There are very few leaf color mutations that are of the nucleo-cytoplasmic interaction type [17].

At present, the research on leaf colors is focused on genetic analysis [11, 18–20]. This method has been currently used in the study of environment stress on plants, regulation mechanism of leaf color, etc. [21–23]. Till date, there is no study reporting the use of proteomics in understanding sesame leaves color, but this method is a powerful approach to identify and isolate different proteins.

In this study, we generated a yellow color sesame mutant, *Siyl-1*, using EMS mutagenesis. The mutant exhibited a stably inherited yellow leaf trait in different years and growing environment. Furthermore, the mutant selfing will separate out three different color types: light-yellow (lethal), yellow-green and normal green.

In order to systematically analyze and parse the mutant molecular regulation mechanism, we proposed to investigate the cytology, genetics, and proteomics of the sesame mutant using the above material. The objectives of the present study were: (1) to clarify the genetic background of *Siyl-1*; (2) to explore the cytological characteristics of *Siyl-1*; and (3) to analyze the protein expression profiles of *Siyl-1* and identify specific protein candidates related with leaf color mutation. This study for the first time revealed the genetic, cytological, and proteomic differences between the leaf color mutant *Siyl-1* and the wild type. This study also provided a foundation for further sesame proteomics research.

## Materials and Methods

### Plant materials

The yellow leaf mutant *Siyl-1* was used for leaf color trait analysis (Fig. 1). The mutant was induced from var. ‘*Yuzhi 11*’ using EMS mutagenesis by the Henan Sesame Research Center (HSRC), Henan Academy of Agricultural Sciences (HAAS), China. Another three sesame varieties with normal leaf color, including *Yuzhi 4*, *Zhengtaizhi 1*, and *Zhengheizhi 1* were used for genetics and cytological analysis of mutation through reciprocal cross with *Siyl-1* (Table 1). Before performing cross hybridization, all these varieties were purified for more than seven generations. All the materials are available from the sesame germplasm reservoir of HSRC, HAAS. In this study, normal green (*yy*) was compared with light-yellow (*YY*) and yellow-green (*Yy*) of *Siyl-1*.

**Fig. 1.**
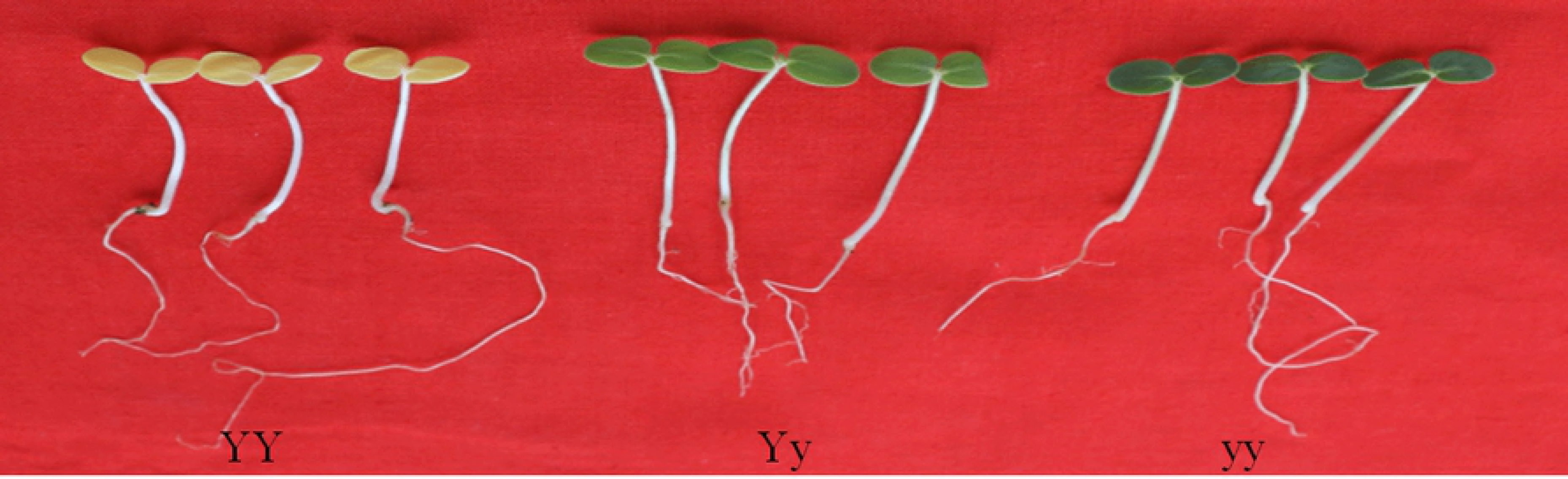
Three phenotypes of the yellow leaf mutant *Siyl-1* in sesame.

**Table 1.** Genetic analyses of the yellow leaf trait in *Siyl-1*.

| Cross combination<br>(Female × Male) | Number of F<sub>1</sub> | Phenotype separation | | | | $\chi^2$ | P |
| --- | --- | --- | --- | --- | --- | --- | --- |
|  |  | Dead with pale | Yellow green | Green | Expected |  |  |
|  |  | yellow leaf<br>(YY) | leaf<br>(Yy) | leaf<br>(yy) | ratio |  |  |
| *Siyl-1* (Yy) × *Siyl-1* (Yy) | 4607 | 1151 | 2328 | 1128 | 1:2:1 | 0.751 | < 0.05 |
| *Siyl-1* (Yy) × Yuzhi 4 (yy) | 59 | 0 | 28 | 31 | 1:1 | 0.068 | < 0.05 |
| Yuzhi 4 (yy) × *Siyl-1* (Yy) | 92 | 0 | 44 | 48 | 1:1 | 0.098 | < 0.05 |
| *Siyl-1* (Yy) × Zhengtaizhi 1 (yy) | 245 | 0 | 119 | 126 | 1:1 | 0.147 | < 0.05 |
| Zhengtaizhi 1 (yy) × *Siyl-1* (Yy) | 192 | 0 | 89 | 103 | 1:1 | 0.880 | < 0.05 |
| *Siyl-1* (Yy) × Zhengheizhi 1 (yy) | 185 | 0 | 102 | 83 | 1:1 | 1.751 | < 0.05 |
| Zhengheizhi 1 (yy) × *Siyl-1* (Yy) | 286 | 0 | 138 | 148 | 1:1 | 0.283 | < 0.05 |

To explore the genetic characteristics of the leaf color trait in the mutant *Siyl-1*, the self-pollution and the cross hybridization of 6 groups were performed using the above materials during 2014-2016 (Table 1). The leaf color trait in each sample was observed three times after germination. Chi-square tests (P = 0.05) were used to determine the segregation significance for the leaf color characteristic.

To perform proteomics analysis, the healthy seeds of the three genotypes of the mutant *Siyl-1* progeny were cultured in artificial growth chambers (GPJ-400, Changzhou, China) at 30 °C with 70% relative humidity and 14 h light/10 h dark cycle for 1 week. Three biological replicates were set. After germination, 3 g of cotyledons per genotype were collected and frozen with liquid nitrogen for subsequent protein extraction. In addition, another 3 g of cotyledons per genotype were collected for chlorophyll content measurement.

### Chlorophyll content measurement

Before measuring the chlorophyll content, 0.5 g cotyledons of three genotypes (*yy*, *Yy*, *YY*) were ground individually using calcium carbonate powder and silica sand, and the chlorophyll was extracted [24]. The chlorophyll of three plantlets was extracted and measured in 3 biological replicates. Chlorophyll content was measured using UV-VIS spectrophotometer (TU-1810, Beijing, China). The absorbance wave length was set at 665 nm, 649 nm, and 470 nm. The photosynthetic pigment content was calculated using the following formulas:

Chlorophyll a content (mg·g^-1^) = (13.95×A_665_-6.88×A_649_)×V×N/W/1000. Chlorophyll b content (mg·g^-1^) = (24.96×A_649_-7.32×A_665_)×V×N/W/1000. Carotenoid content (mg·g^-1^) = [1000×A_470_-2.05×(13.95×A_665_-6.88×A_649_)-114.8×(24.96×A_649_-7.32×A_665_)]×V×N/W/1000/245, where V is the extracted volume (mL), N is the dilution factor, and W is the fresh sample weight.

Total chlorophyll content (mg·g^-1^) = Chlorophyll a content + Chlorophyll b content. Duncan’s multiple-range test was applied to analyze the chlorophyll content.

### Cytological observation

After germination, the 2 × 2 mm sections of young leaves of three genotypes (*yy*, *Yy*, *YY*) were collected and immediately pre-fixed in 3% (pH 7.2) pre-cooling glutaraldehyde fixative. Samples were rinsed with 0.1 mol· L^-1^ phosphate buffer (pH 7.2), fixed in 1% osmium acid solution, and then rinsed using the same buffer. All the fixed samples were dehydrated, impregnated, embedded, and polymerized with the epoxy resin Epon 812. Sectioning was performed using the Power Tome-XL ultrathin microtome, and the samples were double-stained with uranyl acetate and citrate. The chloroplast structure was observed and imaged under a Hitachi H-7650 transmission electron microscope (Hitachi, Tokyo, Japan). For each sample, three biological replicates were observed.

### Protein extraction and 2-DE

Total proteins in leaves were extracted using trichloroacetic acid (TCA)/acetone mixture [25]. Optical density of the protein samples was measured at 595 nm using the Bradford’s method [26]. Samples were then stored at −70 °C for 2-DE electrophoresis. Three biological replicates per sample were performed.

2-DE was performed using the methods of Ma [27]. The 100 mL samples were reserved with 400 μL hydrated sample buffer. SDS-PAGE was performed following the method of Sui [28]. Polyacrylamide gels were stained, washed, and then scanned with a Molecular Imager Pharos FX System (Bio-Rad) (AB Company, Milwaukee, USA). Protein in-gel digestion and MALDI-TOF-TOF/MS analysis

Target protein spots were individually cut from the gel and washed three times with ultra-sterilized water. Each protein sample was de-stained with 50 μL of ammonium bicarbonate (100 mmol·L^-^1 NH_4_HCO_3_). After the excess solution was discarded, 100% acetonitrile was added for 5 min. Then, 5 μL of ammonium bicarbonate solution (50 mmol·L-1 NH_4_HCO_3_) with 10 ng of trypsin was added per sample tube. Each tube was then covered with 20 μL of ammonium bicarbonate solution and digested at 37 °C for 16 h. Finally, the 5 μL of 5% trifluoroacetic acid was added per tube and left for 10 min to terminate the reaction.

After digestion, 1 μL of supernatant was placed on the sample target point. The 1 μL of α-Cyano-4-hydroxycinnamic acid matrix solution was placed on the same point. The target sample was then subjected to matrix-assisted laser desorption/ionization-time of flight (MALDI-TOF-TOF/MS). Mass spectrometry parameters were set as follows: primary MS molecular weight ranges from 700-3500 Da, and secondary MS molecular weight ranges from 40-1050 Da.

### Protein identification and data analysis

Incorporating MS and secondary MS data, raw data files were converted to *.mgf files. The file was submitted to the MASCOT search engine for protein retrieval (http://www.matrixscience.com) (series parameters listed in Table 2). The NCBI nr (non-redundant) database was used for collection of protein information with a taxonomy parameter set to green plants. The function of DEPs was interpreted by using the UniProt database (http://www.uniprot.org). To better understand the biological function and interaction of DEPs, a protein-protein interaction network (PPI) was predicted using the online analysis tool STRING 9.0 (http://string-db.org).

**Table 2.** Parameter information for protein retrieval from MASCOT database

| Options | Setup parameter |
| --- | --- |
| Enzyme | Trypsin |
| Maximum allowable shear loc | 1 |
| Fixed modifications | Carbamido methylation |
| Variable modifications | Oxidized |
| Primary mass spectrometry tolerance | 150 ppm |
| Secondary mass spectrometry tolerance | 0.6 Da |
| P value | $\leq 0.05$ |

### RNA extraction and cDNA reverse transcription

Total RNA was extracted from sesame leaves of all three phenotypes according to the manufacturer’s instructions of an RNA extraction and purification kit (Sangon Biotech, Shanghai, China). The RNA concentration and quality was assessed using a nucleic acid and protein analysis system Nanodrop 2000 (Thermo scientific, Wilmington, USA). The total RNA was reverse transcribed into cDNA with a RevertAid First Strand cDNA Synthesis Kit (Thermo scientific, Vilnius, Lithuania).

### Quantitative real-time PCR analysis

To analyze the gene expression of these DEPs, we explored the gene sequences for these proteins from the genome database of Yuzhi 11 (data unpublished), and performed quantitative real-time PCR (qRT-PCR) analysis. qRT-PCR was performed with FastStart Essential DNA Green Master (Roche, Mannheim, Germany) using the Realplex 4 MasterCycler real-time PCR System (Eppendorf AG, Hamburg, Germany). Each reaction mixture contained 10.0 μL of 2X master mix, 7.4 μL of double-distilled water, 2 μL of diluted cDNA, 0.3 μL of forward primer and 0.3 μL of reverse primer. The qRT-PCR was conducted in accordance with the manufacturer’s instruction. The reaction conditions were as follows: 1 cycle of 50 °C for 2 min; 1 cycle of 95 °C for 10 min; 40 cycles of 95 °C for 15 s; 40 cycles of 60 °C for 20 s; and 40 cycles at 72 °C for 20 s. Three independent biological replicates per sample were performed and three technical replicates were analyzed for each RNA sample. *SiTUB* was used as an internal reference gene to normalize the relative gene expression [29]. The relative gene expression was calculated using Pfaffl method [30]. The primer sequences of the genes of DEPs for qRT-PCR are shown in Table 3.

**Table 3.**
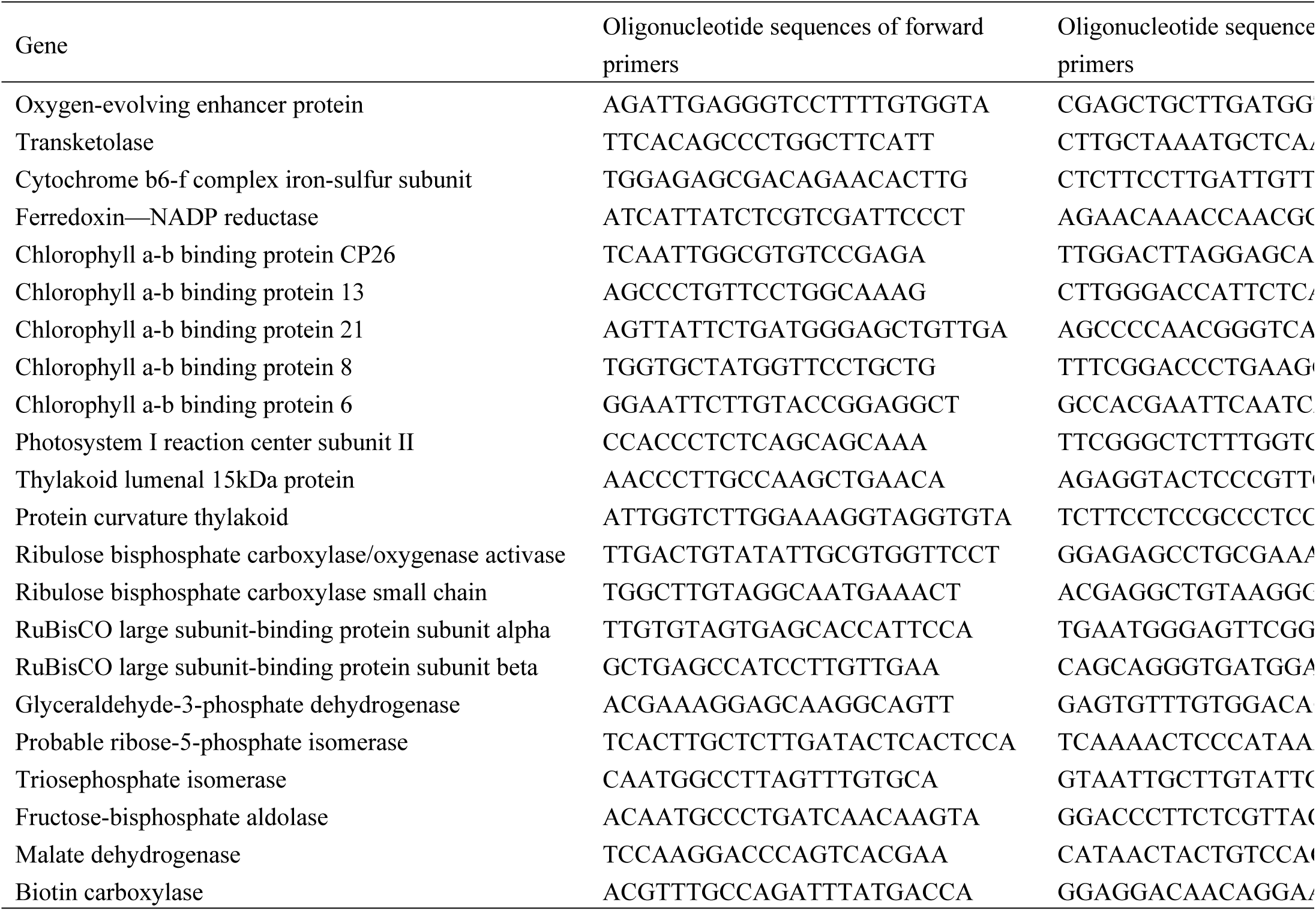
Primer sequences used for qRT-PCR of genes involved in photosynthesis and energy metabolism in *Siyl-1*

## Results

### Genetic analysis of the leaf color mutant Siyl-1

To clarify the genetic background of yellow leaf color trait in mutant *Siyl-1*, we investigated the phenotypes of the self-pollinated progeny and 6 test F_1_ progeny from reciprocal cross between the mutant type (*Yy*) and 3 wild type (*yy*) of *Siyl-1* (Fig. 1 and Table 1). There were three phenotypes in self-pollinated progeny, i.e., light-yellow (*YY*), yellow-green (*Yy*), and normal green (*yy*) with the expected separation ratio of 1 (*YY*): 2 (*Yy*): 1 (*yy*) using χ^2^ tests. The *Yy* with yellow-green leaf could complete the whole life cycle. However, the *YY* always die after 1day of emergence. For the *Siyl-1* (*Yy*) reciprocal crosses, the phenotype separation ratio was consistent with the expected ratio of 1 (*Yy*):1 (*yy*) by χ^2^ tests. The genetic analysis results proved that the mutant *Siyl-1* (*Yy*) is a heterozygosis type, and the yellow leaf trait is controlled by an incompletely dominant nuclear gene; named *Siyl-1* in sesame.

### Leaf chlorophyll content and cell ultra-structure variation in Siyl-1

To understand the cytogenetic mechanism of early lethality of the homozygous seedlings (*YY*) in *Siyl-1* progeny, we first analyzed the variations in the chlorophyll composition of the three genotypes at cotyledon stage (Table 4). Compared with *yy* (0.51 mg·g^-1^), the chlorophyll content in the seedlings of *Yy* and *YY* significantly reduced to 0.29 mg·g^-1^ and 0.01 mg·g^-1^, respectively. The content of chlorophyll b was 0.18 mg·g^-1^ (*yy)*, 0.08 mg·g^-1^ (*Yy),* and 0.02 mg·g^-1^ (*YY*). The total chlorophyll content was 0.68 mg·g^-1^ (*yy)*, 0.37 mg·g^-1^ (*Yy),* and 0.03 mg·g^-1^ (*YY*). The individual contents of chlorophyll a, chlorophyll b, and total chlorophyll also decreased significantly. However, the reduction in the carotenoid content was insignificant in the *Yy* type (0.09 mg·g^-1^) compared to the *yy* (0.10 mg·g^-1^).

**Table 4.** Chlorophyll content of *Siyl-1* progeny at cotyledon stage

| Genotype | Chlorophyll a content<br>(mg·g<sup>-1</sup>) | Chlorophyll b content<br>(mg·g<sup>-1</sup>) | Carotenoid content<br>(mg·g<sup>-1</sup>) | Total chlorophyll content<br>(mg·g<sup>-1</sup>) |
| --- | --- | --- | --- | --- |
| *YY* | 0.01±0.00 c | 0.02±0.00 c | 0.05±0.00 b | 0.03±0.00 c |
| *Yy* | 0.29±0.01 b | 0.08±0.00 b | 0.09±0.01 a | 0.37±0.01 b |
| *yy* | 0.51±0.00 a | 0.18±0.01 a | 0.10±0.00 a | 0.68±0.01 a |

Note: a, b, and c within the columns indicate the significant differences at $P < 0.01$ .

Subsequently, we compared the ultra-structure of leaf cells in the three genotypes (Fig. 2). In *YY*, there were no obvious difference in the location of chloroplast, but their shape changed (Fig. 2A and 2B). The shape of chloroplast changed from a convex lens like shape to a circular type. The lamellar structure was unclear and disordered as a loose strip. The thylakoid volume dropped and the osmiophilic granules increased. In contrast, chloroplasts in *Yy* type was larger and had a thicker spindle shape with a cavity. Meanwhile, gaps were observed in the stacking and folding lamellar structure. The chloroplast structure was abnormal and the osmiophilic granules appeared (Fig. 2C and 2D). As for *yy* progeny (used as control), chloroplasts exhibited a normal spindle shape without cavities (Fig. 2E and 2F). The lamellar structure was clear and folded tightly. Starch grains were absent; while a small amount of osmiophilic granules were present in cells (Fig. 2E and 2F).

**Fig. 2.**
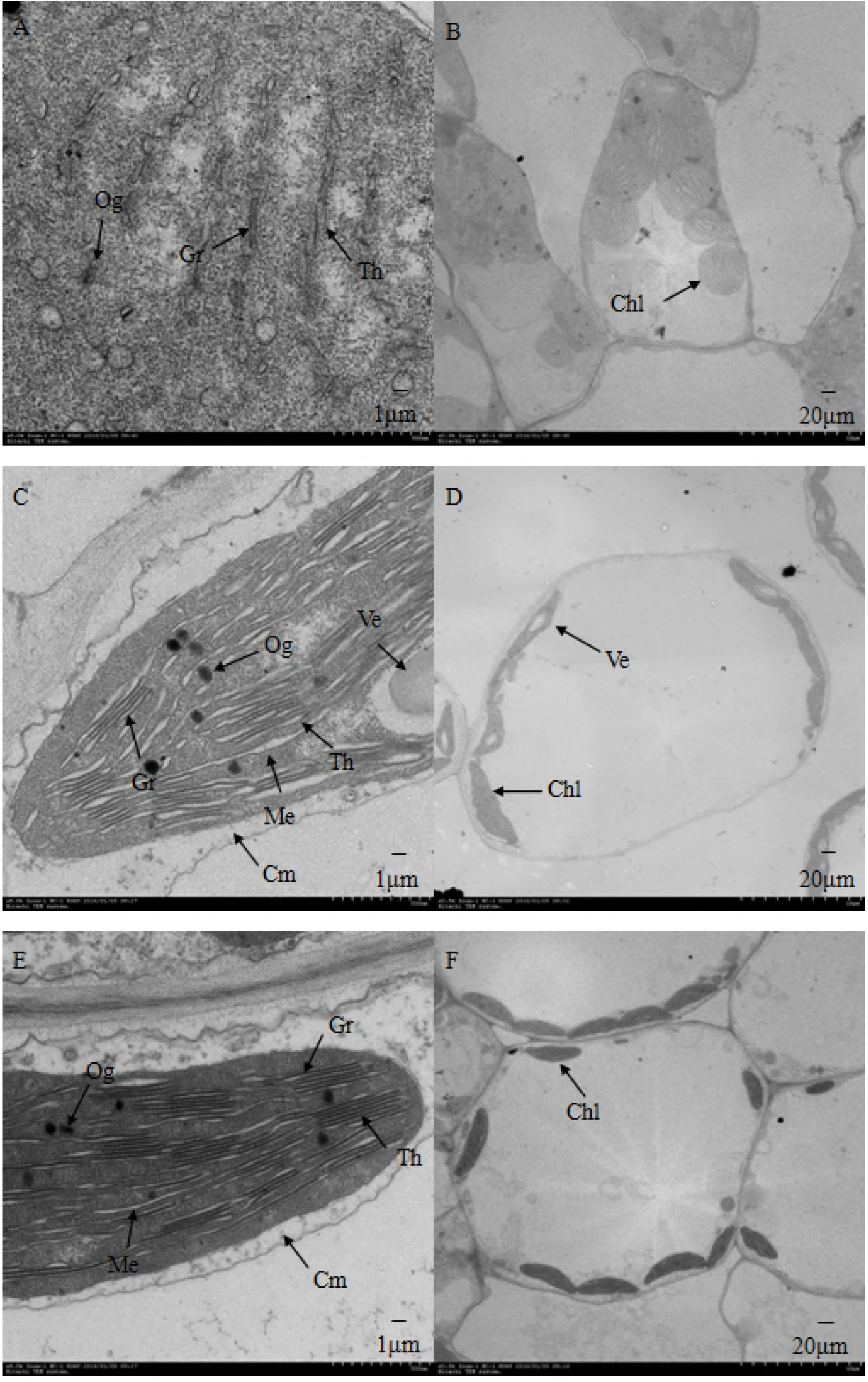
Morphological comparison of chloroplast cells of three genotypes of *Siyl-1*. **A** and **B**: *YY*. **C** and **D**: *Yy*. **E** and **F**: *yy*. Cm: chloroplast outer membrane; Th: thylakoid; Og: osmiophilic globule; Gr: grana; Me: mesenchyme; Ve: vesicle; Chl: chloroplast.

### Comparative proteomics of the mutant type and the wild type of Siyl-1

To further explore the regulation mechanism of sesame leaf color, we performed 2-DE experiment and compared the proteomic differences between the wild type (*yy*), and the mutant type (*YY* and *Yy*) of *Siyl-1* (Table 5). The results showed that 518, 521, and 535 protein spots were obtained from the individuals of *YY, Yy* and *yy*, respectively. The isoelectric point of the protein spots mainly aggregated within a pH range from 4- 7, and their molecular weight generally ranged from 10 to 110 kDa. Protein Gel images of light-yellow (*YY*), yellow-green (*Yy*) and normal green (*yy*) phenotypes were analyzed in PDQuest 7.2 (Fig. 3). A total of 98 individual protein spots with 2-fold up- or down-regulated variation were identified by comparing the two mutants with the wild type (Fig. 4). In these proteins, the expression abundance of 57 proteins changed among *YY/yy*, and 17 among *Yy/yy*. The 98 protein spots were digested with trypsin and analyzed by MALDI-TOF-TOF analysis. Then, these proteins were ultimately screened using the NCBI nr database (Fig. 3). The 98 DEPs are listed in Supplemental Table 1, with spot number, NCBI accession number, protein name, protein score, SC, MP, mutant /wild type, theoretical TMr/Tpl, and experimental EMr/Epl. The Kyoto Encyclopedia of Genes and Genomes (KEGG, http://www.kegg.jp/kegg/pathway.html) analysis grouped the 98 proteins into five functional categories: photosynthesis and energy metabolism (82.7%), synthesis, folding and proteolysis (9.2%), detoxification and antioxidation (5.1%), respiration (2.0%), and defense-related protein (1.0%) (Fig. 5). The mutation of leaf color trait had a tight relationship with the photosynthesis bioprocess in sesame. Among DEPs, there were some down-regulated light-reaction proteins, including oxygen-evolving enhancer protein (OEE), cytochrome b6-f complex (Cyt b6-f), iron-sulfur subunit, ferredoxin-NADP reductase (FNR), chlorophyll a-b binding protein (LHC), and photosystem I reaction center (PS I) (Supplemental Table 1).

**Fig. 3.**
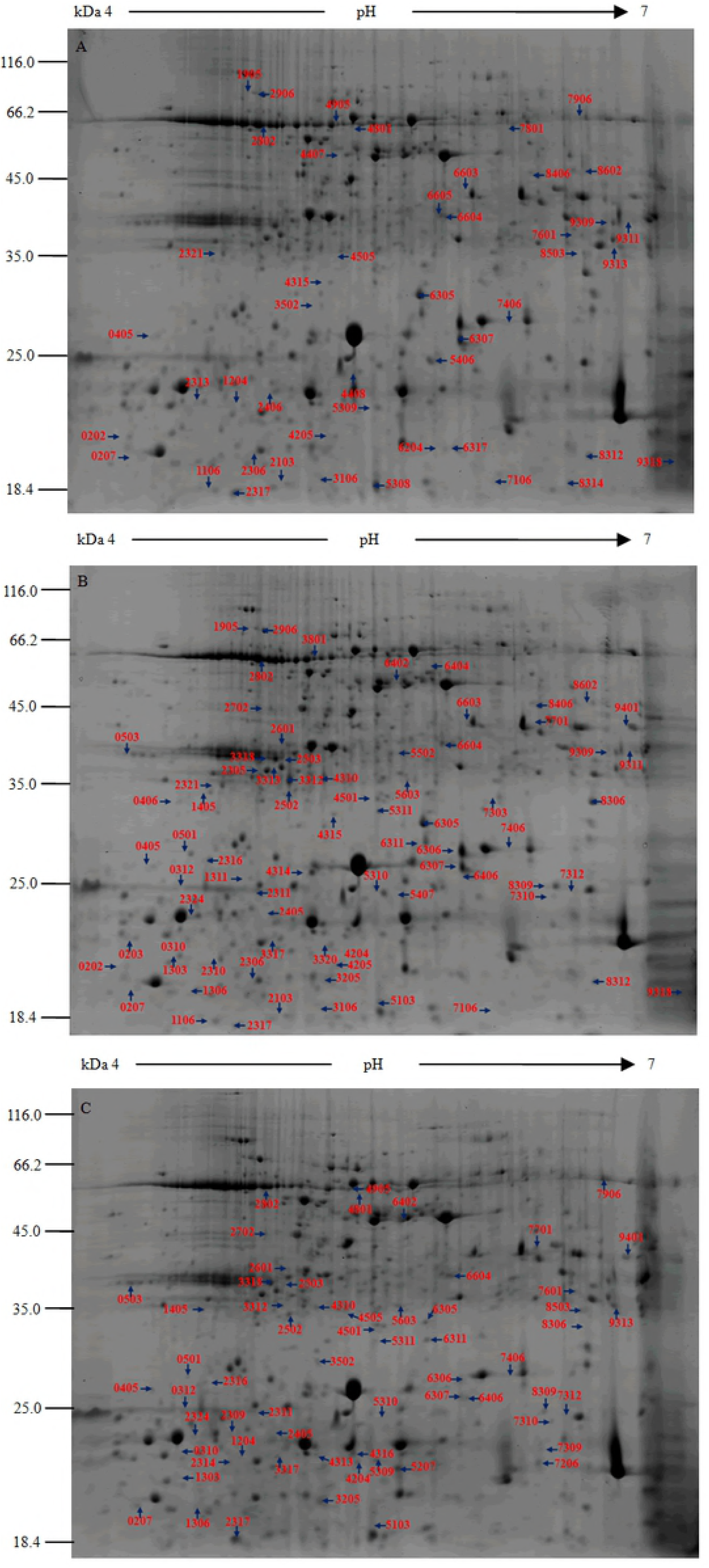
Identification of 98 protein spots according 2DE and MALDI-TOF-TOF MS analysis. The proteins are extracted from the leaves of Siyl-1 during seedling stage. The number signed in the images indicates DEPs. **A**: *YY*; **B**: *Yy*; **C**: *yy*.

**Fig. 4.**
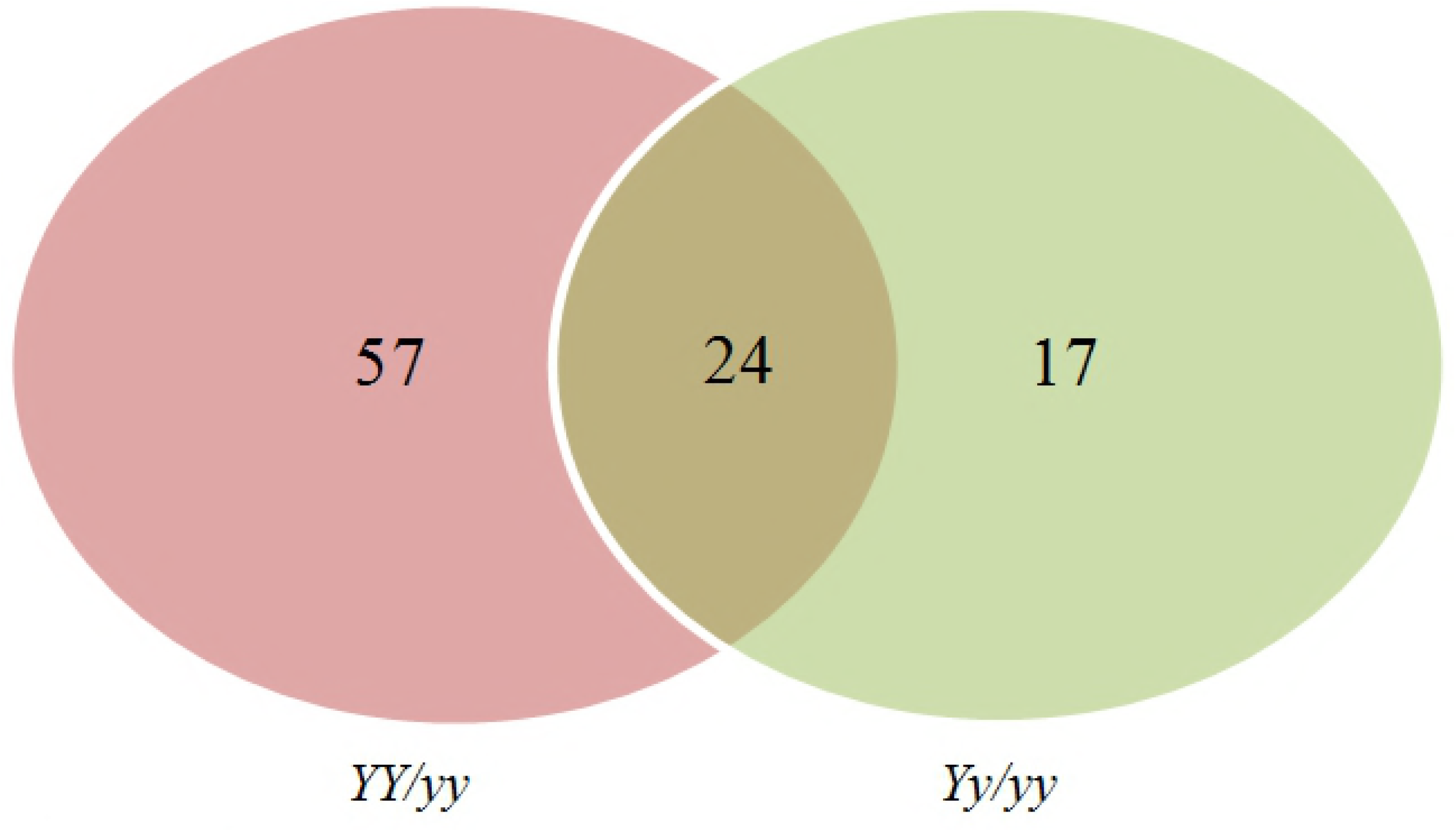
The Venn diagram of the comparing the two mutants with wild type.

**Fig. 5.**
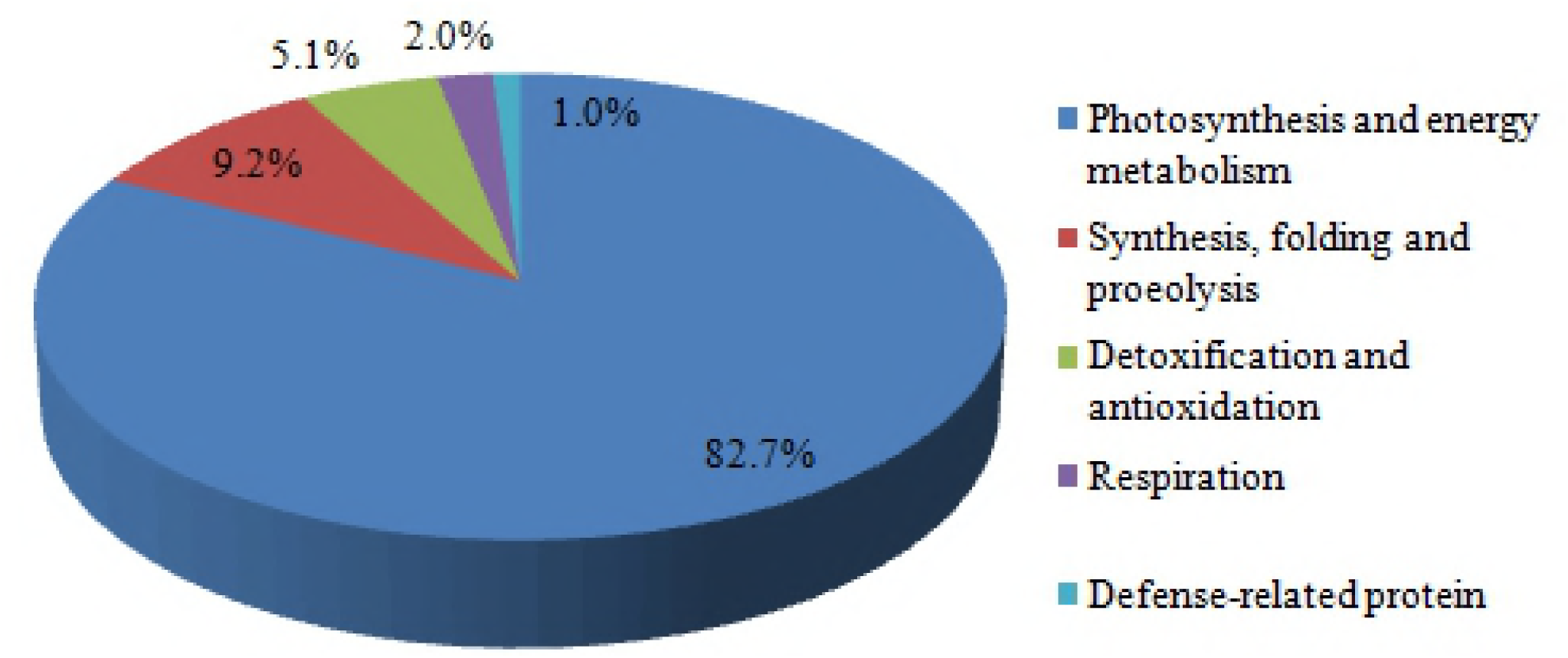
Functional groups of DEPs identified from *Siyl-1*. This classification is based on KEGG (http://www.kegg.jp/kegg/pathway.html) and the literature.

**Table 5.** Proteomics comparison of the mutant types (*YY* and *Yy*) and the wild type (*yy*) of *Siyl-1* using 2D gels

| Item | *YY* | *Yy* | *yy* |
| --- | --- | --- | --- |
| Protein quality (mg) | 4.5 | 4.5 | 4.5 |
| Number of protein spots | 518±15 | 521±15 | 535±15 |
| Number of different protein spots (%) | 74 (14.3) | 87 (16.7) | 47 (8.8) |
| Number of up-accumulated spots | 60 | 63 | 31 |
| Number of down-accumulated spots | 14 | 24 | 16 |

### Abundance patterns of DEPs in Siyl-1

Two mutant types (*YY* and *Yy*) were compared with the wild type (*yy*); the results demonstrated that the expression patterns of 98 DEPs were significantly different. The heat map provides an overview of these DEPs (Fig. 6). In *YY/yy*, the number of down- and up-regulated proteins was 62 and 19, respectively, whereas in *Yy/yy* it was 16 and 25 respectively. More importantly, among these DEPs, there were 11 common DEPs that were down-regulated synchronously in *YY/yy* and *Yy/yy.* Additionally, another 11 common DEPs were up-regulated synchronously in *YY/yy* and *Yy/yy* (Supplemental Table 1). In this study, about half of the DEPs had different molecular mass or isoelectric point. For instance, PSBP1 possessed four different molecular mass or isoelectric point (Supplemental Table 1).

**Fig. 6.**
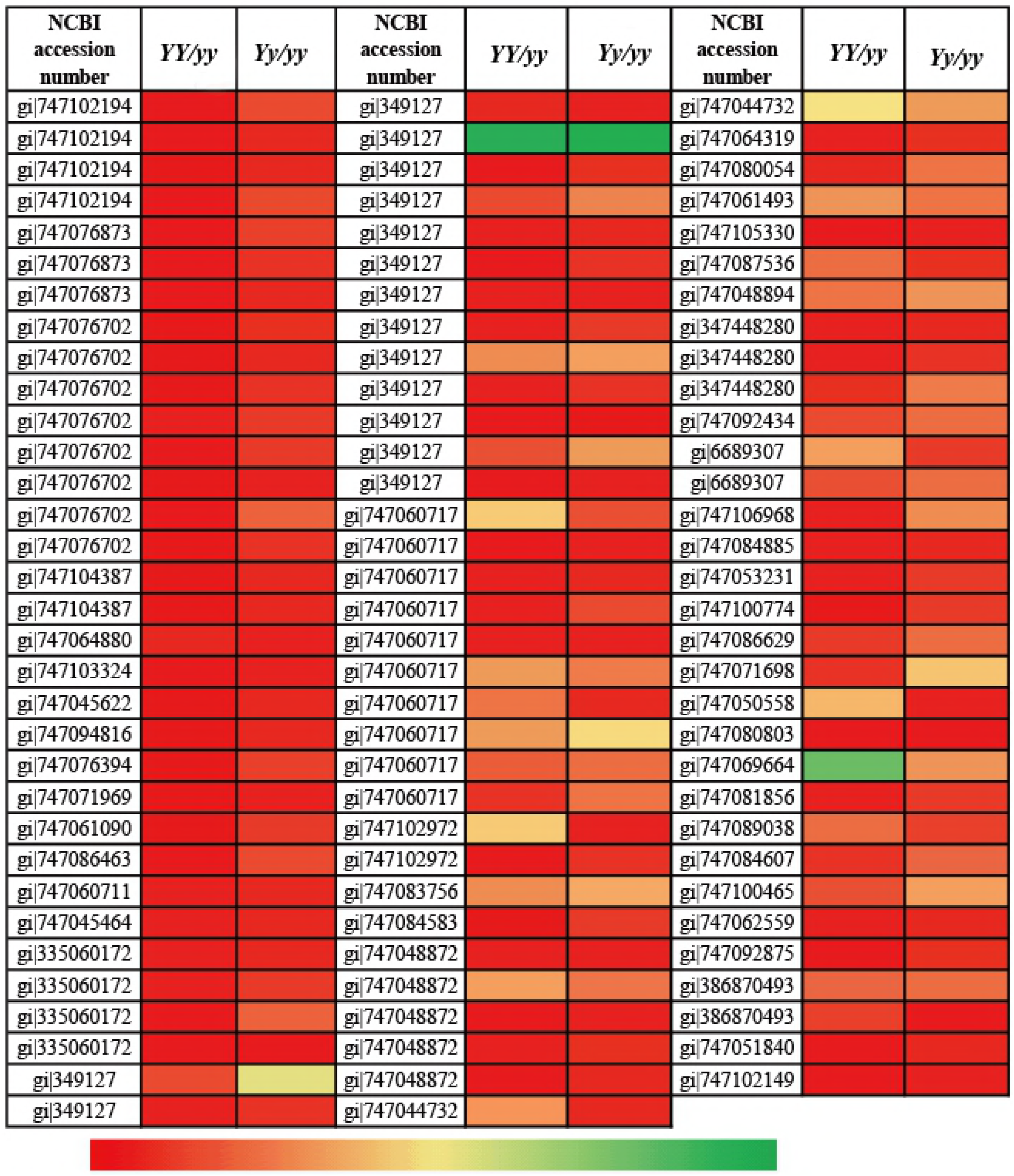
Heat map representation of 98 DEPs identified of *Siyl-1*. Color scores were normalized by the log_2_ transformed of the protein abundance ration of each spot. Red represented increased protein abundance, and green represented decreased protein abundance, respectively.

### Interaction of DEPs

STRING 9.0 was used to analyze the protein-protein interaction and molecular function of DEPs (Supplemental Fig. 1, 2). The results revealed that these proteins formed a complicated interaction network, and most of the core interacting proteins were related with photosynthesis and energy metabolism (Supplemental Table 2). In addition, there are 21 proteins (ATPQ, PB, LHCA1, LHCA3, LHCA5, GS2, RPS1, FBA2, PSBO2, PSI-P, PETC, GAPA, PSAD-2, RBCL, PSBP-1, CYP38, cpHsc70-1, RPE, TRX-M4, CPN60A, and At5g06290); each of them could interact with more than 15 proteins. For instance, there were 27 different proteins that interact with the TRX-M4 (thioredoxin-M4) protein. Similarly, GAPA also interacts with another 27 proteins.

### Expression profiles of genes involved in DEPs of Siyl-1

In order to study the expression relationship between the protein and gene in the yellow leaf mutant *Siyl-1*, we matched the mRNA sequence with the DEPs using the genome data of sesame in the NCBI database (https://www.ncbi.nlm.nih.gov/). A total of 22 genes were identified involved in photosynthesis and energy metabolism (Fig. 7). The qRT-PCR was used to analyze the expression levels of these genes (Fig. 7). There were 2 different gene expression patterns compared with the protein expression (Supplemental Table 1): (1) A total of 7 (31.8%) genes had the same expression trend as the proteins, such as A: OEE, B: transketolase (TKT), C: Cyt b6-f, G: LHC 21, H: LHC 8, J: photosystem I reaction center subunit II, and K: thylakoid luminal 15 kDa protein; (2) The rest of the genes (15, 68.2%) that had a different trend from the proteins expression levels, included D: FNR reductase, E: LHC CP26, F: LHC 13, P: RuBisCO large subunit-binding protein subunit beta, R: ribose-5-phosphate isomerase, I: LHC 6, L: protein curvature thylakoid, M: ribulose bisphosphate carboxylase/oxygenase activase, N: ribulose bisphosphate carboxylase small chain, O: RuBisCO large subunit-binding protein subunit alpha, Q: glyceraldehyde-3-phosphate dehydrogenase, S triosephosphate isomerase, T: fructose-bisphosphate aldolase, U: malate dehydrogenase, and V: biotin carboxylase.

**Fig. 7.**
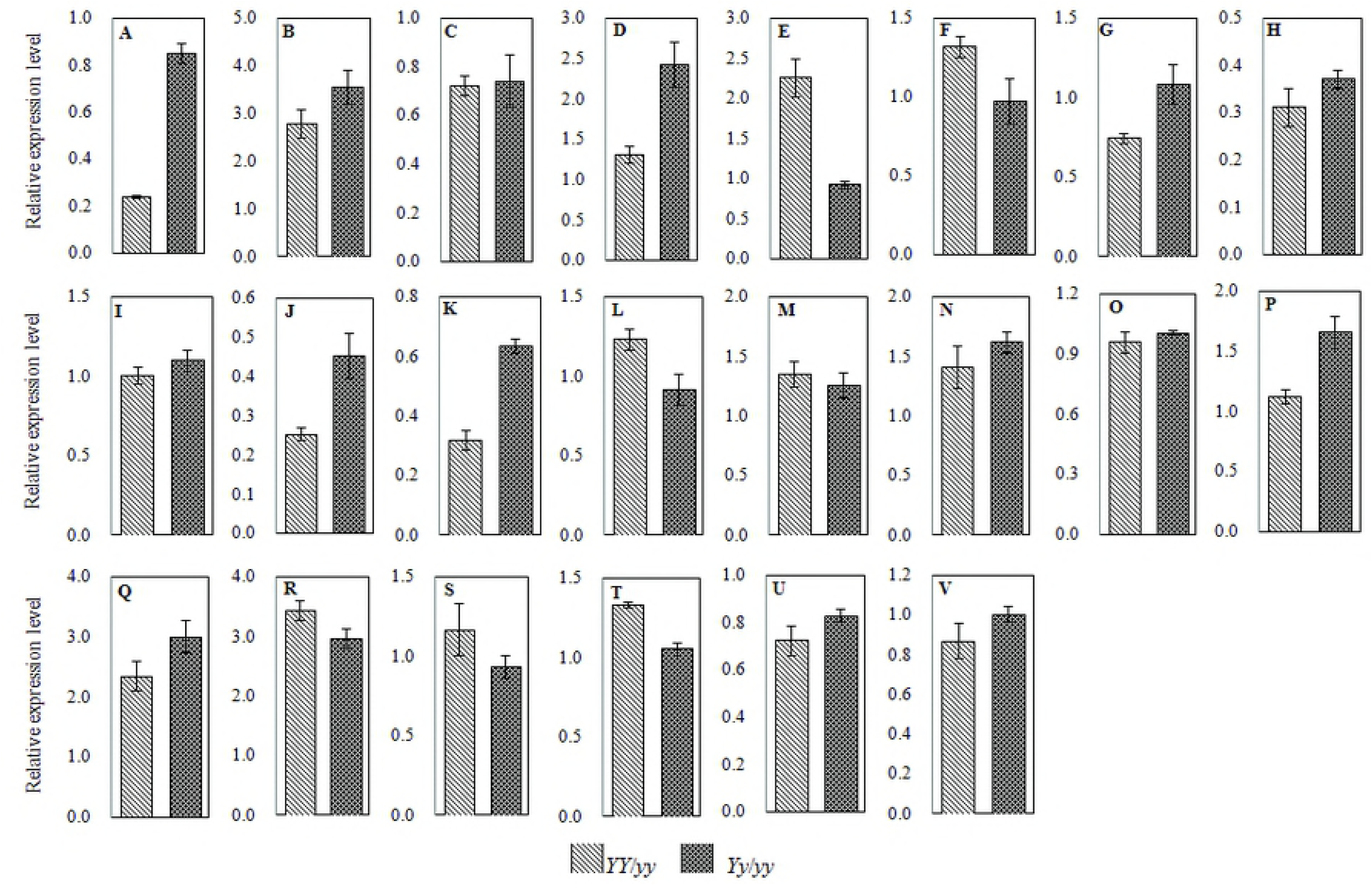
Expression profiles analysis of 22 genes encoding photosynthesis and energy metabolism proteins identified in *Siyl-1*. Standard deviation was expressed by error bars among three independent replicates. A, oxygen-evolving enhancer protein (OEE); B, transketolase (TKT); C, cytochrome b6-f complex iron-sulfur subunit (Cyt b6-f); D, ferredoxin-NADP reductase (FNR); E, chlorophyll a-b binding protein CP26 (LHC CP26); F, chlorophyll a-b binding protein 13 (LHC 13); G, chlorophyll a-b binding protein 21 (LHC 21); H, chlorophyll a-b binding protein 8 (LHC 8); I, chlorophyll a-b binding protein 6 (LHC 6); J, photosystem I reaction center subunit II (PS I); K, thylakoid lumenal 15kDa protein; L, protein curvature thylakoid; M, ribulose bisphosphate carboxylase/oxygenase activase; N, ribulose bisphosphate carboxylase small chain; O, RuBisCO large subunit-binding protein subunit alpha; P, RuBisCO large subunit-binding protein subunit beta; Q, glyceraldehyde-3-phosphate dehydrogenase (GAPA); R, ribose-5-phosphate isomerase (Rib-5-P); S, triosephosphate isomerase (TPI); T, fructose-bisphosphate aldolase (FBA); U, malate dehydrogenase (ME); V, biotin carboxylase.

## Discussion

Leaf color mutation in plants is common and there is a rich variation in nature. The mutated genes not only impact the biosynthesis and degradation of chlorophyll by direct or indirect way, but also affect the growth and development of plants [31]. In this study, for the first time we analyzed the genetic, cytologic, and proteomic data of the leaf color mutant *Siyl-1*.

### Genetic analysis

Several leaf color mutants were found in rice [11], barley [32], soybean [33], maize [34], and wheat [35]. These studies identified that most of the leaf color mutants are controlled by a single recessive nuclear gene. For example, many leaf mutants of soybean[33], wheat [36], and rice [37] were all controlled by a single recessive nuclear gene. In addition, there are a few leaf color mutants, in which the traits are controlled by the cytoplasmic gene. This kind of mutants exhibits cytoplasmic inheritance and displays a mosaic phenomenon in the development process [38].

In our study, we identified that the yellow leaf trait was controlled by a nuclear gene, which displayed incomplete dominance heredity. This will serve as a new resource material for sesame research. Similarly, in the leaf color mutants found in wheat [35], maize [34], barley [39], and tobacco [40], the traits were all controlled by an incompletely dominant gene. And their offspring also had three different phenotypes; light-yellow (*YY*), yellow-green (*Yy*), and normal green (*yy*) leaf color, similar with our study.

### Analysis of pigment content and morphological characteristics

The chloroplast is an essential organelle for photosynthesis in plants, which contains chlorophyll and carotenoid. In the present study, the content of chlorophyll in mutant was significantly lower than the wild type. This is a common phenomenon in leaf color mutants [41]. In general, the morphological characteristics of normal chloroplast are elliptic or approximate elliptic shape, stable size, and number. In the *Siyl-1* mutant, the shape and internal structure of the chloroplast changed dramatically. These abnormal changes indicated that the chloroplast development has been restricted, chlorophyll biosynthesis was abnormal, and leaf color was affected. Similar studies on leaf color mutant also appeared in other crops, for example, chloroplast development was significantly delayed in rice, arabidopsis and wheat leaf color mutants [42–44].

### The function analysis of DEPs

Comparative proteomic analysis provides a qualitative and quantitative expression profiling of differential proteins, and helps to systematically comprehend of the underlying molecular events [11]. In this study, using the proteomics approach, a total of 98 DEPs had been identified, and the different protein expression profiles in the three genotypes of *Siyl-1* progeny revealed that the mutagenesis had an effect on the accumulation and expression of these proteins. In addition, about half DEPs were detected in multiple spots with different molecular mass or isoelectric point, which implied the existence of posttranslational modifications and isoforms [45]. More importantly, most (82.7%) of DEPs were involved in photosynthesis and energy metabolism. Additionally, 63.3% DEPs were down-regulated in *YY/yy* comparison, which were consistent with the variation in the mutated leaf chlorophyll content and cell structure. In the present study, many down-regulated photosynthesis-related proteins (OEE, Cyt b6-f, FNR, LHC and PS I) involved in photosystem I and II were located on the photo membrane and played an important role in photosynthetic electron transport. OEEs are components of photosystem II (PS II) reaction center and promote the activation of oxygen, which can effectively organize the transfer of protons and electrons among the photosynthetic electron transport chain, receptors, and donors [46]. OEE1 is a manganese-stabilizing protein required for PS II core assembly/stability and OEE2 is responsible for catalyzing the splitting of water. Based on our proteomic analysis, both OEE1 (1–7) and OEE2 (8–15) were significantly decreased in *YY* compared to the wild type plants [47, 48]. These data indicated that the ability of transferring protons and electrons was strongly suppressed in *Siyl-1*.

The LHC is an important component of light harvesting complex in the thylakoid membrane, as the protein can capture and transduct the light energy, increasing the light energy capture efficiency and adjusting the light energy distribution [49]. All the chlorophyll-binding proteins decreased in NYC1 like mutant [50]. The chlorina mutant of barley lacks chlorophyll b; the thylakoid membrane polypeptide [51]. When *Stay-Green* is transiently overexpressed in mutant of *arabidopsis*, both chlorophyll *a* and *b* are degraded, and an *NYC1*-deficient mutant did not degrade chlorophyll *b* and LHCII under *SGR* overexpression [50, 52]. In this study, LHCs (20-24) were significantly decreased in *YY* when compared to the wild type plants. Therefore, we speculate that those different proteins are related to the ability of capturing light in *Siyl-1*.

The Cyt b6-f acts as a bridge to PSI (plastocyanin: GAPOR) via electron transfer [53]. The activity of the light-harvesting complex of photosystem II kinase is regulated by a Cyt b6-f component(s), which responds to the balance of electron flow from photosystem II to photosystem I via the plastoquinone pool [54]. Thylakoids could not be phosphorylated under any experimental condition in the Cyt b6-f-less of mutant [55]. FNR located in thylakoid membrane, can catalyze the electron transfer from the iron-sulfur leading to NADP+ [56], when FNR gene is mutated, the chlorophyll level also reduces in the mutant, and exhibits the yellow-green leaf phenotype [57]. In our study, the Cyt b6-f (16, 17), FNR (19), PSI protein (25) and thylakoid protein (26, 27) were significantly decreased in *YY* when compared to the wild type plants. Therefore, we hypothesized that the change of thylakoid structure causes PSI protein deactivation, which caused the loss of electron transfer function of the Cyt b6-f.

TKT is a key enzyme in the Calvin cycle of photosynthesis and the pentose phosphate pathway in all organisms. The low amounts of TKT should repress the efficiency of Calvin cycle and pentose phosphate pathway, hindering the photosynthesis rate and resulting in plant death [58]. Thus, the TKT plays an important role in plant defense and growth. It is interesting that the protein expression level of TKT increased in *YY* when compared to the wild type plants in our study. This mechanism of protein expression of TKT, along with other similar proteins such as biotin carboxylase and triosephosphate isomerase (TPI) warrants further investigation.

Other DEPs involved in the Calvin cycle, energy conversion mechanism, and synthesis, among others had substantial difference in the protein expression profiles; for example, the 1, 5-diphosphate ribulose carboxylase/oxygenase (RuBisCO), glyceraldehyde-3-phosphase dehydrogenase (GAPA), and adenosine triphosphate synthase (ATPase). The physiological and molecular mechanisms of these processes need further investigation.

### Expression levels of genes related to DEPs

Photosynthesis in leaves provides the basis for grain formation in crop production [35]. Leaf color mutants are valuable materials for understanding chloroplast development, chlorophyll biosynthesis and metabolism, photosynthesis, and for identifying the functions of genes [37, 50, 55]. Biosynthesis and expression of genes might be strongly perturbed by mutation [59]. For instance, in a nuclear mutant of *C. reinhardtii*, the expression level of the oxygen-evolving activity were reduced due to the absence of OEE2 polypeptides [60]; a significant increase of TKT in both enzyme activity and transcript level under stress was also observed in mutant of yeast *Candida magnoliae* [61]; FNR does not influence the distribution of excitation energy between photosystem II and photosystem I, and also does not impact the occurrence of ‘light-state transitions’ [62]; in *arabidopsis*, the expression level of LHC was also increased in the *Mybs1* mutant, whereas the *Mybs2* mutant exhibited decreased expression of this gene [63]; the RuBisCO large subunit (RbcL) protein expression levels of the pale-green plants were much higher than that in *Fln2-4* analyzed by qRT-PCR. RbcL protein promote the accumulation of RuBisCO [64]; for *D. salina*, glyceraldehyde-3-phosphate dehydrogenase (*GAPDH*) was reduced to 41.2% and 67.4% in transformants G1 and G2 of the wild-type, respectively [65]. The mutant plant contained DEPs, and differential genes expression levels were evident between the wild type and the mutants. In this report, the expression trend was consistent with previous studies of *OEE* [60] and *TKT* [61]. *GAPDH* was opposite to the results shown in *D. salina* [65]. However, the expression trend was up and down for chlorophyll a/b-binding protein gene [63] and RuBisCO [64]. These results suggest that the greater influence of gene biosynthesis and expression due to mutations may be partly due to different mRNA levels of each gene.

The genetic, cytological, and proteomic analyses of the sesame yellow leaf mutant, *Siyl-1* (*Yy*) were performed in this study. The yellow leaf mutant (*Yy*) is controlled by an incompletely dominant gene. The chlorophyll content dramatically decreased and the ultrastructure of the leaf chloroplasts were significantly altered in *Siyl-1*. For the first time, a proteomic analysis was performed on sesame leaf color mutant. A total of 98 DEPs were identified in *YY* and *Yy* genotypes. All DEPs were classified into the functional groups of which photosynthesis and energy metabolism (82.7%) was predominant. The number of down-regulated proteins was greater in *YY* than in *Yy*. Most of the DEPs involved in photosynthesis and chlorophyll biosynthesis were inhibited in *YY*. Among the identified proteins, OEE, Cyt b6-f, CHL, PS I center and thylakoid protein, involved in morphogenesis of chloroplast, photosynthetic electron transport, and light absorption, were found to be down-regulated in *YY*. The inhibition of these proteins was partly due to the decreased expression levels of their genes. The greater influence of gene biosynthesis and expression due to mutation may be partly due to different mRNA levels of each gene. These findings provide the basis for further exploration of the molecular mechanisms underlying the leaf color mutation in sesame.

## Supporting information

**Supplemental Table 1.** A total of 98 DEPs identified in a yellow leaf mutant *Siyl-1* in sesame. (DOC)

**Supplemental Table 2.** Interprotein function in a yellow leaf mutant *Siyl-1* in sesame. (DOC)

**Supplemental Fig. 1.** Simplified diagram of some key biological processes for photosynthesis and energy metabolism of *Siyl-1*. (TIF)

OEE, oxygen-evolving enhancer protein; Cyt b6-f, cytochrome b6-f complex; LHC, chlorophyll a-b binding protein; FNR, ferredoxin-NADP; Trx, thioredoxin; RuBisCO, ribulose-1,5-bisphosphate carboxylase/oxygenase; GAPA, glyceraldehyde-3-phosphase dehydrogenase; TPI, triosephosphate isomerase; FBA, fructose-bisphosphate aldolase; TKT, transketolase; Rib-5-P, ribose-5-phosphate; GS, glutamine synthetase; PPlase, peptidyl-prolyl cis-trans isomerase; ME, malate dehydrogenase; PS I, photosystem I reaction center.

**Supplemental Fig. 2.** Analysis of a functional network of DEPs using STRING 9.0 (http://string-db.org). (TIF).

The 0.4 is required interaction score of medium confidence of minimum. The different color lines and nodes represent the different relationship types. Line colours represent the types of evidence in predicting the associations. Gene neighborhood (green), gene fusions (red), gene co-occurrence (blue), co-expression (black), experimentally determined (purple), from curated databases (light blue), textmining (yellow) and protein homology (light purple).

## Acknowledgments

This work was financially supported by the [China Agriculture Research System 1] under Grant [CARS-14]; [Plan for Scientific Innovation Talent of Henan Province 2] under Grant [184100510002]; [Key Project of Science and Technology in Henan Province 3] under Grant [151100111200]; [Importing the International agricultural Sciences and Technology Program 4] under Grant [2016-X05]; and [Henan Province Specific Professor Position Program 5] under Grant [SPPP2016].

## Author Contributions

G.T. Performed the experiments and wrote the manuscript draft. W.S. and T.Y.guided the experiments and performed the data analysis. C. J., L.C. and W.L. participated in the proteomic data analysis. W.Y., L.F., Z.Y. and W.D. performed the experiments. Z.H. conceived of the experiments, revised the manuscript and determined the final version.

## Conflicts of Interest

The authors declare no conflicts of interest.

